# A feature selection strategy for gene expression time series experiments with hidden Markov models

**DOI:** 10.1101/392761

**Authors:** Roberto A. Cárdenas-Ovando, Edith A. Fernández-Figueroa, Héctor A. Rueda-Zárate, Julieta Noguez, Claudia Rangel-Escarenõ

## Abstract

Studies conducted in time series could be far more informative than those questioning at a specific moment in time. However, when it comes to genomic data, time points are sparse creating the need for a constant search for methods capable of extracting information out of experiments of this kind. We propose a feature selection algorithm embedded in a hidden Markov model applied to gene expression time course data on either single or even multiple biological conditions. For the latter, in a simple case-control study features or genes are selected under the assumption of no change over time for the control samples, while the case group must have at least one change. The proposed model reduces the feature space according to a two-state hidden Markov model. The two states define change/no-change in gene expression. Features are ranked in consonance with three scores: number of changes across time, magnitude of such changes and quality of replicates as a measure of how much they deviate from the mean. An important highlight is that this strategy overcomes the few samples limitation, common in genomic experiments through a process of data transformation and rearrangement. To prove this method, our strategy was applied to three publicly available data sets. Results show that feature domain is reduced to up to 90% leaving only few but relevant features yet with findings consistent to those previously reported. Moreover, our strategy proved to be robust, stable and working on studies where sample size is an issue otherwise. Hence, even with two biological replicates and/or three time points our method proves to work well.

## Introduction

High dimensional genomic data is described by many features, however, many are either redundant or irrelevant. Identifying those is key in order to claim results are trustworthy. Data mining techniques, machine learning algorithms or statistic models are applied to classify features but at the cost of other important problems such as model over-fitting or the increase of computational resources and higher analysis cost [1] [2]. A possible approach to this classification problem is to reduce data dimensionality with feature extraction (FE) or with feature selection techniques (FS) [3] [4].

FS is a technique often used in domains where there are many features and comparatively few samples, particularly used in microarray or RNAseq studies where there are thousands of features and a small number of samples [5] [6]. FS is the process of eliminating irrelevant features in the data by extracting only a subset of informative ones. Its main objectives include avoid overfitting, eliminate noise in the data, reduce algorithmic rate of convergence, improve model performance and, give a more accurate interpretation of features in the data [7]. It is important not to be confounded with FE such as principal components analysis or compression, where they use the projection of the original set to a new feature space of lower dimensionality. In FE, the new feature space is a linear or even non-linear combination of the original features [4]. FS on the other hand, identifies relevant features without altering the original domain.

Due to the high dimensionality of most gene expression analyses, it is necessary to select the most relevant features to get better results interpretation and a deeper insight into the underlying process that generated the data. However, the noisy data and small sample size pose a great challenge for many modelling problems in bioinformatics making necessary to use adequate evaluation criteria or stable and robust FS models [7].

In general, FS techniques can be classified into three main categories: filters, wrappers, and embedded [7] [8]. Filters take as input all the features and reduce them into a relevant subset independent of the model parameters. Wrappers select a subset of features using a search algorithm, then estimate the model parameters for that subset and perform an evaluation test for each model. Wrappers use FS for model identification by selecting the model parameters that best fit the training data and has the highest evaluation score. The embedded approach takes all the features at once, maximizes a learning algorithm optimizing model performance and outputs both the reduced feature set along with its model parameters. In this work we address the classification of time series gene expression data using two embedded processes feature selection and hidden Markov models.

A variety of FS techniques that have been proposed can be classified into parametric [9] and non-parametric methods [10]. The non-parametric methods aim to make a less stringent distribution assumptions, however, validation in the context of small sample sizes is a challenge. Parametric methods assume there is a given distribution for the observed data. The most common parametric latent variable models are the Gaussian mixture models (GMM) and hidden Markov models (HMM). The mixture model is often used to model multimodal data, while the HMM is often used for modeling time series data [8].

In most applications of HMMs, features are pre-selected based on domain knowledge and the feature selection procedure is completely omitted. Some methods have been explored to reduce the feature space by using HMMs as stated in Adams and Beling [11]. However, FS strategies specifically with HMM are sparse. For example, the work of Zhu *et al.* [12] shows a wrapper FS approach to get the best model and then the feature subset for a continuous HMM. The authors proposed a new set of continuous variables, defined as salient features, to avoid searching the space of all feature subsets and to prevent losing information about the original variable. The salient features have proved their effectiveness for FS in GMM [13]. Finally, they apply a variational Bayesian framework to infer the salient features, the number of hidden states and the model parameters simultaneously.

Adams *et al* [11] propose a feature saliency hidden Markov model. This model also uses feature saliencies variables and they represent the probability that a feature is relevant by distinguishing between state-dependent and state-independent distributions. If the number of hidden states is known, this approach simultaneously provides maximum a posteriori estimates and select the relevant feature subset by using the expectation-maximization algorithm. Finally, the most recent work is introduced in Zheng *et al* [14]. Their strategy combines a hidden Markov model, a localized feature saliency measure and two t-Student distributions to describe the relevant and non-relevant features, to accurately model emission parameters for each hidden state. All the parameters are estimated using a Variational Bayes framework.

Most of these methods use saliency parameters additionally to those required by the model, therefore, when analyzing genomic data the increase of variables to be estimated from data becomes a hurdle. Besides, the number of hidden states necessary to model the data also affects the total parameters to estimate. Hence, a strategy that make use of a minimal number of parameters to get the most relevant features from a data set is indispensable to study genomic data.

In this article we present a novel strategy for FS capable of selecting and ranking relevant genes based on the changes between conditions and successive times, using an embedded FS technique with a HMM at its core. Lack of replication is handled using a novel data rearrangement that overcome the limitations of few samples in genomics experiment designs. To prove its efficiency, the strategy is applied to three publicly available data sets.

## Results

### Overview of the strategy

We present a strategy that selects the most relevant features (genes) from high dimensional genomic experiments with longitudinal design for one or multiple conditions. The approach first compares gene profiles over time on the affected samples against those in the control group. Next, it scores the relevant features providing a ranking that can be used to further reduce the data set.

The complete pipeline takes an expression matrix from either a microarray or an RNAseq experiment as input and returns the list of ranked features. The main steps in the process involve transforming and rearranging input data, estimate the model parameters, evaluate time course expression profiles and select relevant features providing a ranking score for such genes. Code for each step in the pipeline is structured as a collection of single functions that allow user to customize methods, the full pipeline is available as an R-library. Some plots are also included. Fig 1 shows the schematic representation of the proposed strategy. Source code can be found as supplementary information S1 File.

**Fig 1.**
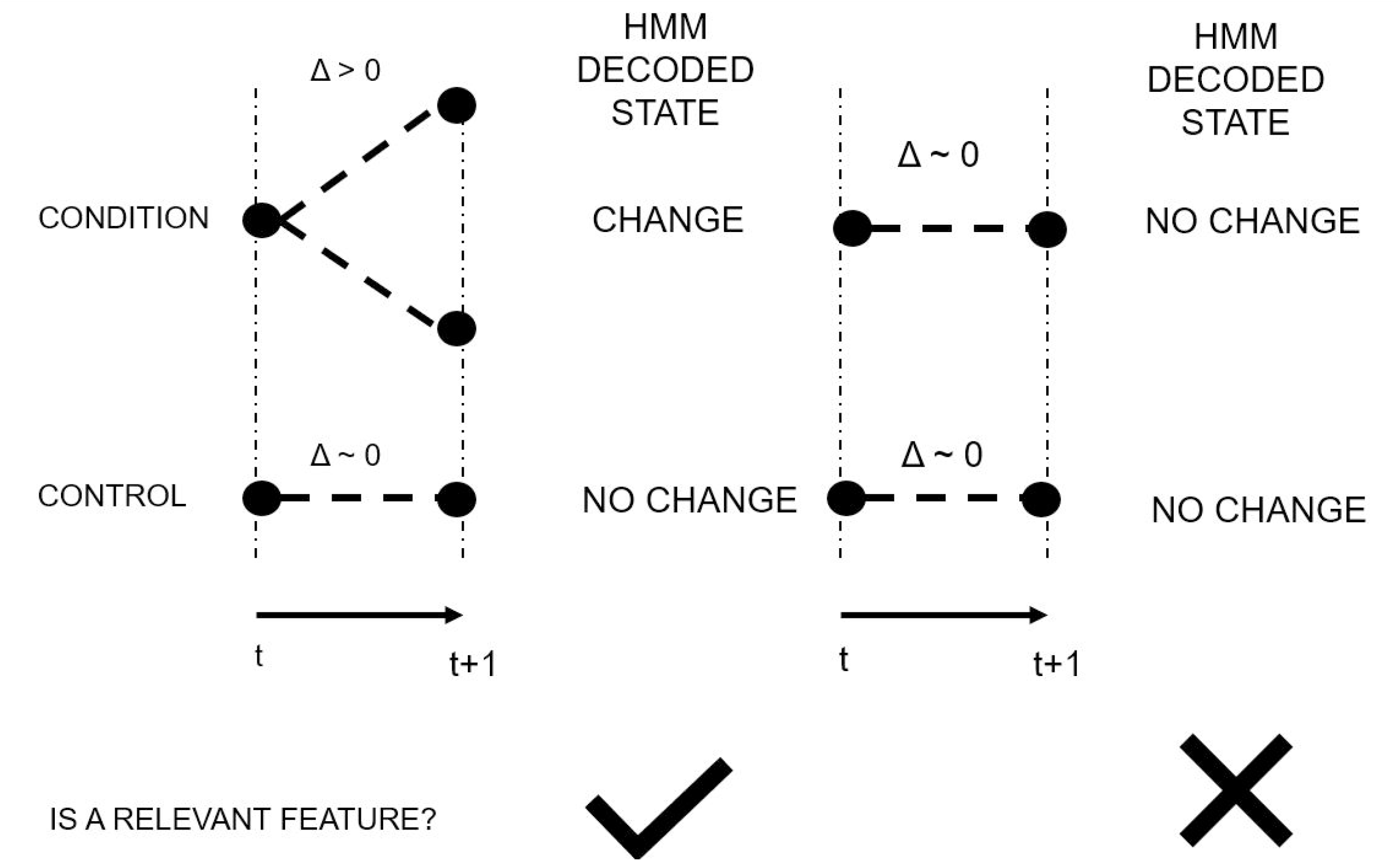
Feature Selection strategy pipeline. The expression matrix is preprocessed to estimate the HMM parameters. With the fitted model, the features are evaluated and compared to filter out the expression profiles with a flat behavior. Finally, the selected features are scored and ranked to give a better interpretation and a deeper insight into the underlying process that generated the data.

A key element of this strategy is the use of the Occam’s razor or law of parsimony in the state transition model used for the HMM. The hidden state complexity is reduced to a minimum with a two-node clique that is capable of fully describing system dynamics by only considering a change or no change in gene expression over time. The ‘change” state would model either up or down regulation. Therefore, with only two states we are able to define three different behaviors.

### Validation using real data

The strategy was applied to three different publicly available datasets, two from the Gene Expression Omnibus [15] and one from the Japanese Toxicogenomics Project (TGP) [16]. We used RNA-seq data (GSE75417) comprised of 6 time points, 2 conditions and 3 replicates [17]. Illumina Microarray gene expression data (GSE39549) consisting of 9 time points, 2 conditions and 3 replicates [18]. Affymetrix Microarrays (TGP) a variety of studies on hepatotoxic compounds made up of 3 time-points, 1 control, 3 conditions and 2-to-3 replicates. These datasets were uploaded as .Rdata to be used with the R-package of our method at github (https://github.com/robalecarova/FSHMM).

#### Ikaros induced B3 cells

The first example, based in the work by Ferreiros *et al* [19] where B3 pre-B-cell line is transduced with mouse stem cell virus retroviral vectors encoding wild-type Ikaros or Aiolos tagged with a hemagglutinin epitope followed by an internal ribosomal entry site and GFP. B3 cells containing inducible Ikaros were treated with 4-hydroxy-tamoxifen and sampled at 0, 2, 6, 12, 18 and 24 hours after Ikaros induction. The control counterpart (not induced) were also sampled at the same times. Data are available at GEO with accession number GSE75417. Gene expression data comprised 2 conditions, 6 time points per condition and 3 biological replicates per time-point leading to 36 RNAseq samples.

Results were compared to those reported in Ferreiros *et al.* [19] which, provided a list of differentially expressed genes as well as the enriched pathways in their Excel Supp 1. Only two comparisons were selected mainly because we only had access to those results. Further comparison included an enrichment analysis with DAVID [20] to match the approach used in what was originally reported. According to the GO terms, results showed many shared pathways, although the number of genes in each differed. We found that 44% of pathways are shared by the two approaches the remainder 56% involves leukocyte cell-cell adhesion, its regulation and some immune system cascades such as JAK/STAT. The cut-off value used, was a natural p-value of 0.05. The list of common and exclusive pathways are available as Supplementary data S1 Table.

We should consider though that the FSHMM strategy used all samples and all time-points to train the model which, presents a better idea of the dynamics in the experiment as opposed to analyzing isolated time-points. Details of genes and pathways are available as Supplementary data S2 Table and S3 Table.

#### High-fat diet in mouse model

A second example with longitudinal design is the one published by Kwon *et al.* [18] and available through GSE39549. It is on an *in vivo* mouse model with two conditions high-fat and normal diet. Both with same time measurements, nine time points at 0, 2, 4, 6, 8, 12, 16, 20 and 24 weeks of high-fat or normal diet intake. The authors analyzed differential expression between both diets. However, they focused on diet effect at each time without considering the longitudinal nature of the study or a temporal dependency. They reported a total of 2037 differentially expressed genes as a result of adding all same-time contrasts. Using FSHMM the number of relevant genes obtained were 1922. We could not report level of agreement between the two studies because the list of differentially expressed genes was not released with their paper. However, a gene set enrichment analysis was performed on the FSHMM results using DAVID, see Supplementary data S4 Table. A caveat regarding this analysis is that search in DAVID involves a variety of parameters associated to different databases, the authors did not elaborate on which ones were used. Therefore, we decided to use the *GOTERM BP FAT* considering the cellular process to which they belong.

Their results showed an enrichment analysis that favors pathways involved in immune response, metabolic process and response to wounding. As stated before, these enriched pathways only considered the treatment *vs* control contrasts. When we compared these results with those obtained by FSHMM, common pathways are those associated to immune response. Hence, either time-by-time comparisons or considering the whole system the level of correlation is good. However, when proposing a time course experiment we should also consider studying the system dynamics instead of partitioning as if we had multiple pair-wise studies. Details of this analysis are available in S4 Table, the shared pathways are shown in yellow, comparing them with Table 2 in Kwon *et al* [18]. In terms of genes found, the authors report Emr1, CCL2, 6 and 7, Adam8, IL1rn, Itgam, CD3, 4, 9, 14 and 180, TLR1, 3, 6 and 7, Tgfb1, Irf5, Mmp12, Col1a, Col2, Col3a1, Col4a5, Col8a1, Col9a3, Col16a1, Ctsa, Ctsb, Ctsk, Ctsl, Ctss and Ctsz as some of the genes differentially expressed in the comparison between same-time contrasts. FSHMM classified as relevant CCL2, 6 and 7, CD14 and CD180, TLR1, 6 and 7, Irf5, Mmp12, Col3a1 and Col1a2. Leading to over 50% of genes found in common. On the other hand, the authors report that the Resistin signaling is activated through the NF-*κ*B transcription factor. In the enriched pathways found with FSHMM, the I-*κ*B kinase/NF-*κ*B signalling was reported, an important finding that coincides between the two analyses.

#### Toxicogenomics data

The third and final example was a dataset from a time series experimental design to study hepatotoxic compounds [16]. The selected compound was Carbon Tetrachloride (CCl_4_) as it is known to be one of the most potent hepatotoxins [21] and is usually used in scientific research to investigate possible beneficial effects from hepatoprotective agents [22] [23]. The study involves only three time measurements at 2, 8 and 24 hours after dose administration with control and three doses: none, low and middle with two replicates leading to a 18 samples. This is an *in vitro* experiment limited in terms of observation space and it was chosen precisely because it represented the *worst case scenario* when it comes to a longitudinal study. It represented the most challenging data to evalulate FSHMM on. Interestingly enough, the number of selected features yielded 1878 genes. The list of all of them is reported in S5 Table as Supplementary information. Pathway analysis using DAVID led to enriched GO terms mainly related to cell regulation, cell and nuclear division, adherens juntion and cell-cell signaling. This correlated to the toxic behavoir of CCl_4_ in liver. Nevertheless, to have another perspective of the results, the KEGG database [24] was used with pathways selected as relevant if they had a p-value lower than 0.05. The significant pathways were also consistent with DAVID and were related to cell regulation, nuclear division, adherens junction and cell-cell signaling. However some of the most relevant pathways that were statisticaly significant were Apoptosis, RNA degradation and the most important was the Carbon metabolism as this is directly related to CCl_4_. When classification parameters were tighten to get more over-represented pathways using the top 200 features, the Nucleotide excision repair pathway was enriched. This pathway is a mechanism to recognize and repair bulky DNA damage caused by xenobiotic factors as compounds, environmental carcinogens, and exposure to UV-light. All of this correlates to what is known about CCl_4_ and its toxicity [21]. Aquiring this level of knowledge with so limited information made FSHMM a perfect choice even on studies this short. The enriched GO terms and details are in Supp S6 Table while the KEGG pathways are in Supp S7 Table.

In summary, for all three cases the feature space was reduced and almost 90% of the variables were filtered out, Table 1. Remarkably, the limited size of replicated observations and time-points were not an issue for the FSHMM to reduce the feature space to a more manageable and yet informative size. The presented method makes use of all time measurements of replicated gene expression values as input and is able to get the most relevant genes by using a HMM as feature classifier.

**Table 1.**
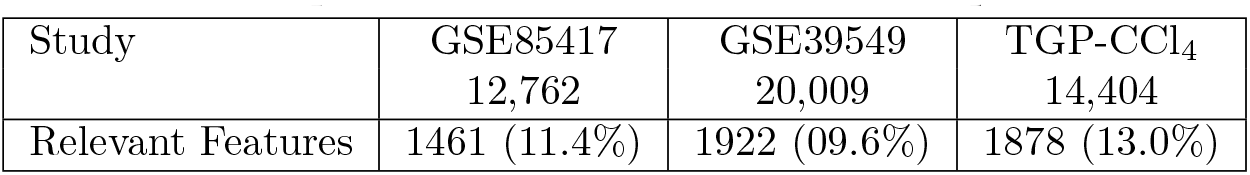
Number and percentage of relevant features per dataset.

|  |  |  |  |
| --- | --- | --- | --- |
| Study | GSE85417<br>12,762 | GSE39549<br>20,009 | TGP-CCl<sub>4</sub><br>14,404 |
| Relevant Features | 1461 (11.4%) | 1922 (9.6%) | 1878 (13.0%) |

## Discussion and Conclusions

Results using the FSHMM strategy outputs a subset of relevant features which is relatively low compared to the original set but yet, it proved to be informative even in situations where number of replicates is as low as two and time series only involves 3 point measurements. The three datasets used to validate our method were chosen because they covered microarray and RNAseq data. Also, because they offered opportunity to compare its efficiency at a pathway level as it was the case of Ikaros data, or gene-to-gene level as we did for the high-fat diet study or on common knowledge after literature review on results from the toxicogenomics data. In all three cases, the strategy efficiently reduced the number of relevant features, simplified the analysis, maintain the time series nature of the studies and provided an insight to the system dynamics. Features were selected according to three scores. One evaluates gene perturbation over time adding more value to genes that fluctuate more over time. A second score evaluates the magnitude of those perturbations and the third score evaluate statistical significance by means of consistency among replicates.

The parameters of the proposed HMM estimated from all three data sets had the same mean vector in their emitted normal distributions. This happens because the HMM models the magnitude of the change in expression instead of the actual expression level. This fact agrees with the hypothesis that only a small set of genes change, and according to our assumptions only those changing in the condition group with no changes among the control set are considered. Therefore, the distributions are centered around zero and this further reduces the number of parameters to estimate from data. Parameters estimated from data using FSHMM strategy were compared to the models presented in [8] [12] and [14], with the conclusion that saliency variables were found not to be necessary, making it all more feasible for a genomic context where the number of observations is low so the less parameters to estimate, the better. The FSHMM used two hidden states to represent relevant features and non-relevant features similar to Zheng *et al*, however the multivariate normal distributions proposed let the user to provide all the replicates with its inherent variability to model the system dynamics.

Also, it is essential to highlight the role of data preprocessing but even more important the input data rearrangement. For our FSHMM strategy, each gene represents a different observation sequence in the training matrix. Therefore, even if the sample size is low, the model parameters can be estimated from data. Moreover, with the feature randomization, it is less probable to overfit. We also analyzed the idea of adding a third hidden state to model the up-regulation, down-regulation, and no-change. Results showed it was not worth it to add complexity by increasing the dimensionality of the hidden vector but instead keep it as simple as possible would allow us to handle studies with minimum counts of data points. The model feeds from changes in expression instead of expression levels themselves, still we can model the sign of the state that emitted the observation. Thus, the two hidden state transition graph is capable of modelling the desired dynamics without increasing the model complexity.

## Methods

### Feature selection with a hidden Markov model

The Feature Selection with a hidden Markov model (FSHMM) strategy starts with an already normalized gene expression matrix, a vector of time points, the biological conditions, the number of replicates and if necessary a set of parameters to customize the model estimation. Data is rearranged into matrices, one per each condition. Then, it is necessary to remove the offset value of each feature by computing the differences in gene expression between consecutive time points. Therefore, instead of using the expression level of each feature for the model parameters, it will receive the change in expression from two consecutive times.

The selected model is a HMM with two hidden states as proposed in Adams *et al* [8] and Zheng *et al* [14]. However, in the proposed model the states represent a non-relevant change in expression (N) and a relevant change (C). The former has the function of a null-hypothesis where most of the features will reside, while the later is the alternative hypothesis. The observations are assumed to be drawn from a multivariate normal distribution with dimensionality equal to the number of biological replicates or a univariate normal distribution if the replicates are summarized. The proposed model is shown in Fig 2. Parameters are estimated using the Expectation-Maximization algorithm and the path of hidden states *X* for each feature per condition is decoded using the Viterbi algorithm. Each feature decoded path is compared to the null-hypothesis that is represented by a sequence of non-relevant changes; if the feature rejects it, then it is selected as relevant. Finally, each relevant feature is ranked by the number of changes, the magnitude of each change and the biological replicates behaviour.

**Fig 2.**
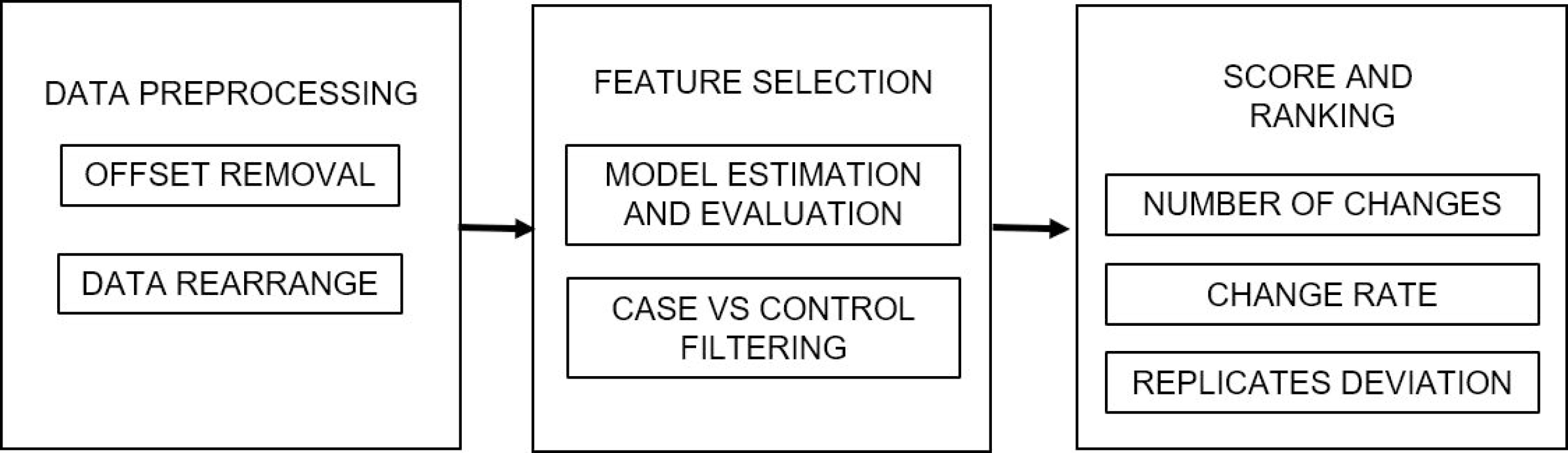
Feature selection hidden Markov model. The state transition graph has two hidden variables: N – Non-relevant expression change and C – relevant expression change. Each state can emit an observation vector with dimensionality equal to the number of biological replicates and it follows a multivariate normal distribution. The model parameters depicted are the transition probabilities and the emission probability of each state.

### Hidden Markov model

Hidden Markov models are stochastic processes based on Markov chains, where the states *X*_1:*T*_ and *X_t_* ∈ *x*_1:*N*_ are connected through a transition matrix. Each state *X_t_* can produce a measurable observation *Y_t_*. These observations only depend of the system’s present state, *P* (*Y_t_|X_t_*) [25] [26]. Unlike the Markov chains, the state variables in the HMM are hidden, this means that they cannot be measured. However, it is assumed that they follow a Markov chain and the transition probability matrix follows a multinomial distribution. If the observations are categorical, then they also follow a multinomial distribution. However, if they are continuous they are assumed to follow a univariate or multivariate normal distribution. Each hidden state has its own mean vector and variance/convariance matrix.

### Decoding - Viterbi algorithm

The decoding function is used to get the hidden states that were traversed by the Markov chain. To decode the states in a hidden Markov model the most commonly used algorithms are the *posterior decoding* and the *Viterbi algorithm* [25] [27]. The Viterbi algorithm is a greedy optimization approach to get the most probable path traversed by the Markov chain. The algorithm computes the best set of hidden states that can explain the present observation, starting from the first one. As it greedily tries to get the best path, it looks for the previous hidden state that maximizes the current joint probability. The algorithm continues to get the best values for each observation until the full sequence has been analyzed. Once it has the most probable outcome, it retraces back its path and outputs it as the most probable path that generated the observation sequence.

### Expectation-maximization algorithm

The Expectation-Maximization (EM) algorithm is applied when there is missing data, or an optimization problem does not have an analytical solution but can be simplified by assuming the existence of hidden variables. The EM algorithm objective is to maximize the complete data set log-likelihood in a two-step procedure. In the first step, it computes the function’s expected value to fill the missing data. And in the second step the algorithm maximizes the model parameters given the *complete data set*. The process is iterated until the convergence criteria are met [28].

### Computational pipeline

The proposed strategy was divided in three stages as stated in Fig 1, each one fulfills a specific objective and are sequentialy executed. These stages are explained below.

#### Data preprocessing

The first step in the pipeline is the data rearrangement and transformation. Each condition is organized in a 3D matrix. Each feature *g* is represented as a matrix with the time points as columns and replicates as rows, Eq 1. The arrangement differs from the usual 2D matrices where the rows contain the features and the columns have time-points and biological replicates. In this strategy an innovative arrangement is proposed, where each condition is organized in a 3D matrix with time points as columns, replicates as rows and as many slices as genes or features are included. With this data arrangement, the few samples limitation that arise in genomic experiments is overcome, given that each feature will be a different observation sequence to fit the model. For a finer parameter estimation, the replicates can be treated as they are or can be summarized by taking the mean or median of them per time measurement.

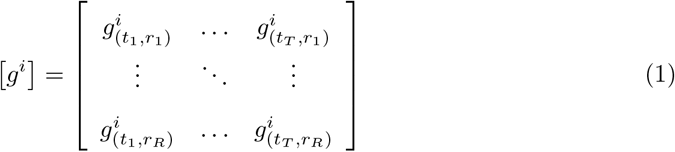

Then, each feature expression value its transformed to get the expression change by substracting the two consecutive time points, Eq 2. With this transformation, the offset value is removed. It is necessary to remove this value because there are features with the same time profile but are shifted given its initial expression value. Thus, without the offset, they can be compared and analyzed as the same profile, Fig 3. However, as result from the transfomation, the observation matrix columns are reduced from size *T* to *T −* 1. For the next step, as it estimates the model parameters, the order of the features may bias the estimation. Therefore, a randomization step is done to shuffle them. This process is based in the preprocess of the sub-sampling cross-validation approach to avoid overfitting [29].

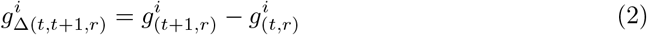

**Fig 3.**
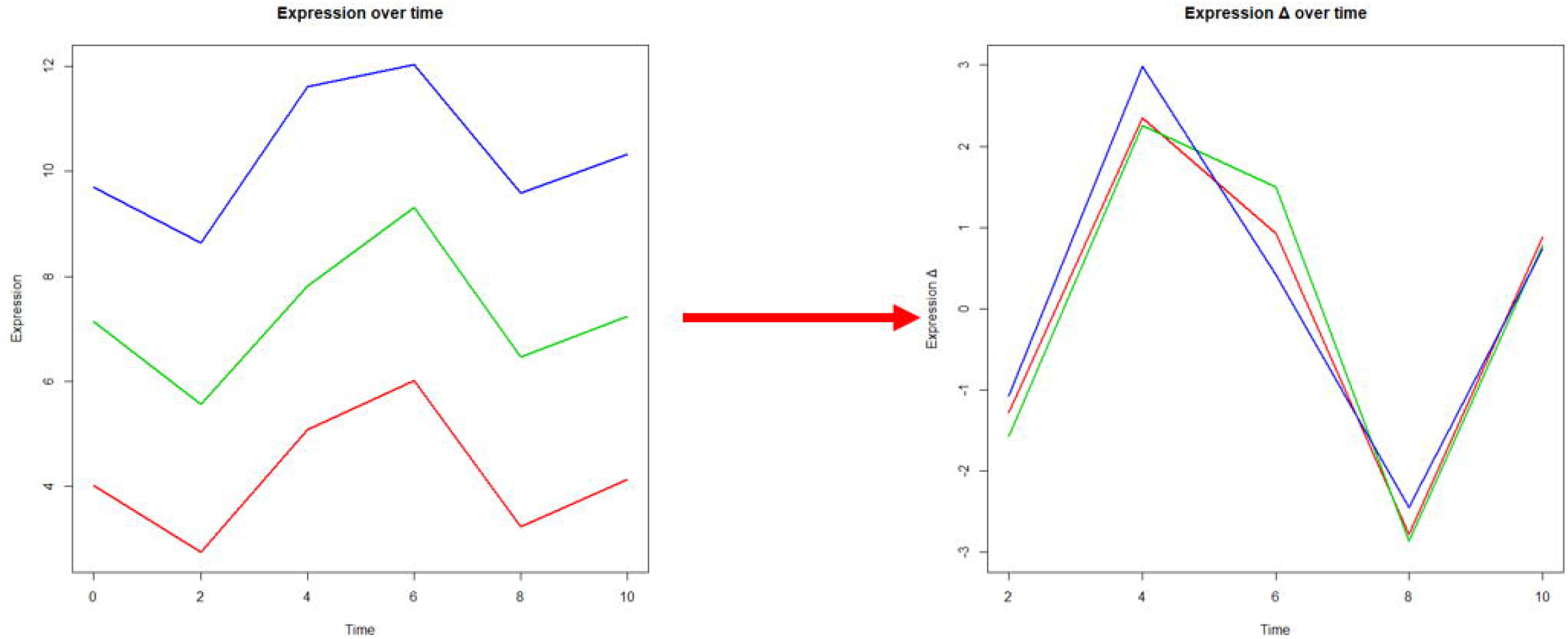
Feature transformation. Different features may have the same time profile, but their initial expression value may shift them. The offset removal makes them comparable and more manageable when analyzing them in the next steps of the proposed strategy.

#### Feature selection

After the data has been organized, the shuffled features are used to estimate the model parameters. With the HMM fitted to the data, the Viterbi algorithm is applied to each feature in each condition, Fig 4. Once each gene has its hidden path decoded, they are compared. In the case of multiple conditions, if at least one condition has a change, but the control is flat, the feature is considered as relevant. And in the case of only one condition without a control or baseline to compare with, all the features that have a flat behavior are discarded, Fig 5.

**Fig 4.**
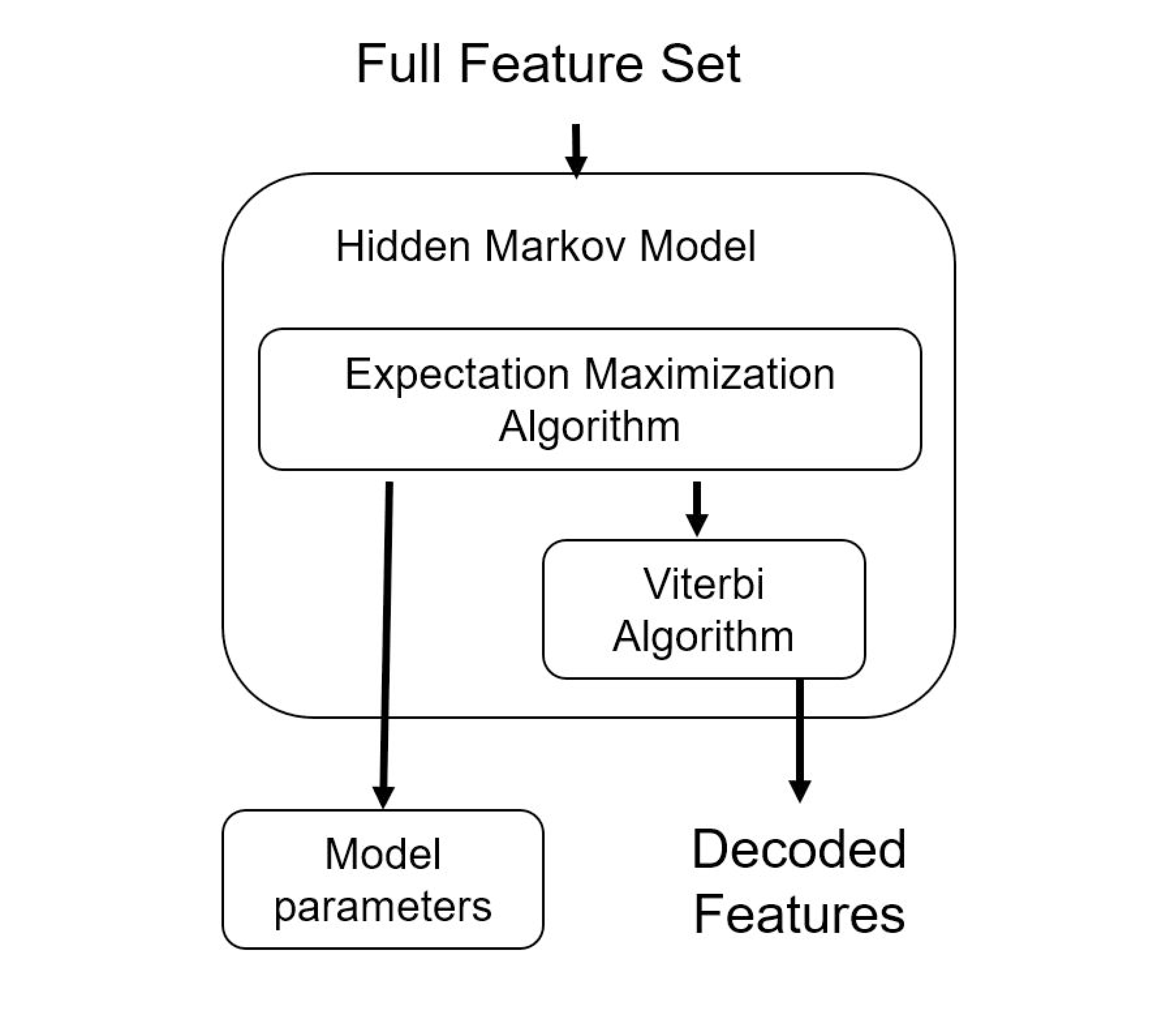
Feature selection embedded technique. The full set of shuffled features is used as the input of the EM algorithm to estimate the HMM parameters. Then the features from the *g*_∆_ matrix are input to the Viterbi Algorithm, and each gene is assigned with the path hidden states traversed by each condition. By comparing he condition’s path, a subset of relevant features is proposed.

**Fig 5.**
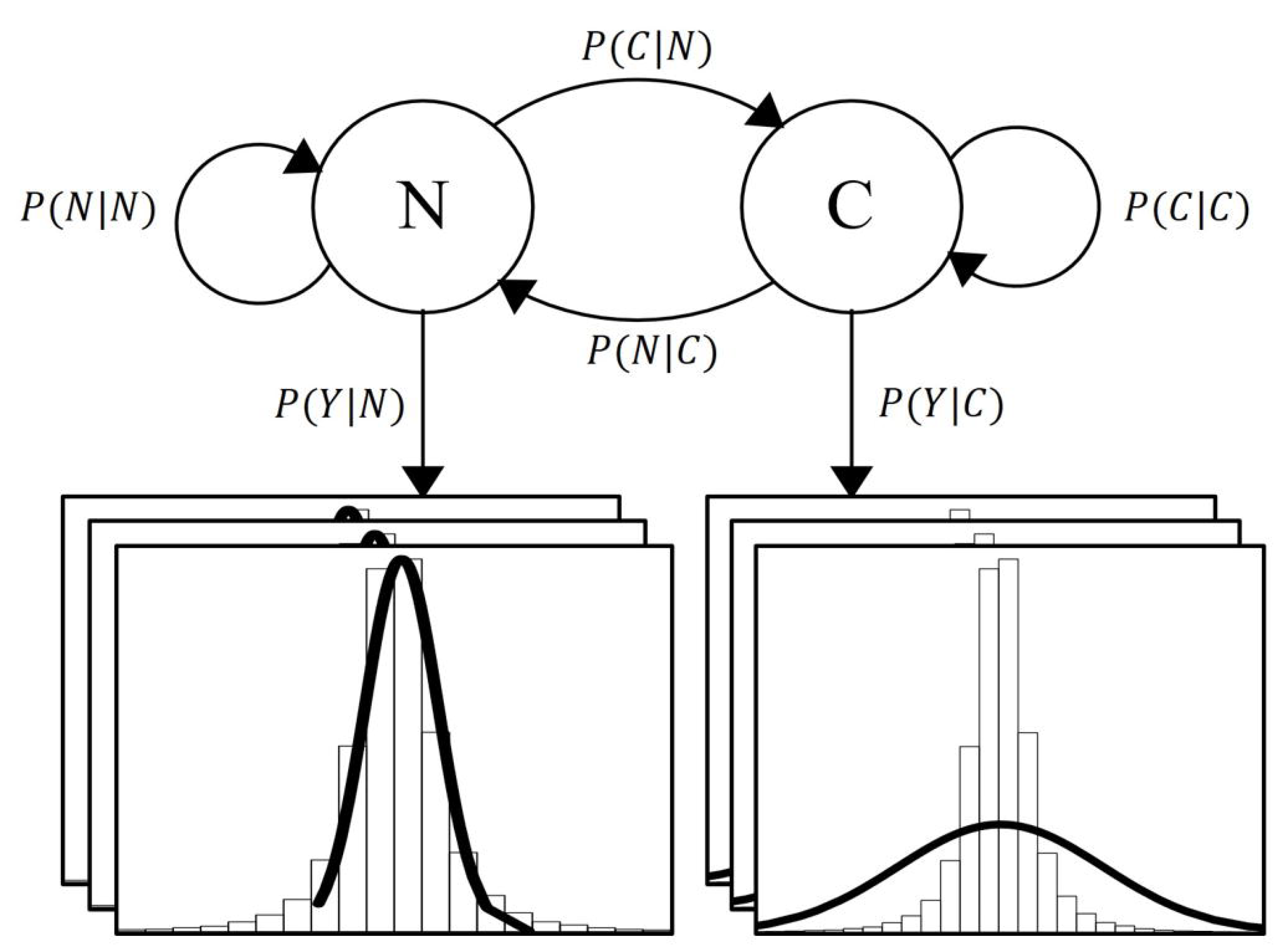
Case-Control hidden state path comparison. If there is a change in expression between consecutive times, then the Viterbi algorithm will set this Δ value in the Change state. When the case is compared, the feature is deemed as relevant only if the control value is decoded as a No Change state.

#### Score and ranking

After the dataset has been filtered. Each relevant variable is evaluated with three different scores:

1. Number of changes across time (#*Ch^i^*). Represents the number of changes that occurred in the time series, in a multiple condition experiment it also considers the feature in each one (*Z*). The greater the number of changes decoded by the Viterbi algorithm (*X* = *C*), the better the score will be, Eq 3.

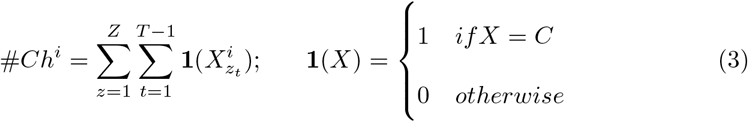
2. Magnitude of change (||∆^*i*^||). It represents how much each relevant variable changes. Even if the feature has only one change in time, if it was very large, this variable will have a good score, Eq 4.

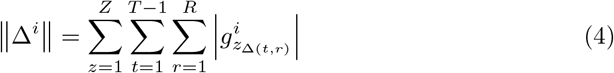
3. Quality of replicates (*scoreR^i^*). It represents the variability between biological replicates. The greater the difference, the lower the value of this score, Eq 5.

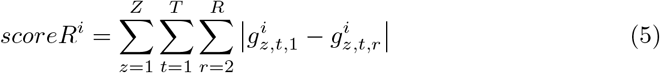

With these scores, variables are ordered and their place in the list represents their rank. The rank serves as filter to find the most important genes within the selected ones.

## Supporting information

**S1 File. FSHMM R-package.** A source code R-package that is ready to be installed. It contains the strategy proposed in this work.

**S1 Table. GO terms found in common with Ferreiros *et al* and exclusive to FSHMM.** Excel book with the enriched GO terms found with FSHMM. The first sheet contains the GO terms in common with the Ferreiros *et al* work, and the second sheet has those found exclusively with the FSHMM strategy.

**S2 Table. Relevant features found in the GSE75417 RNA-seq data.** Table with the gene symbol of the all relevant features found in the GSE75417 RNA-seq data with the FSHMM strategy.

**S3 Table. Enriched GO terms in the GSE75417 RNA-seq data.** Table with the over repressented GO terms using the relevant features found in the GSE75417 RNA-seq data and DAVID. The table has the GO term, its description, the p-value, adjusted p-value, q-value and its relevant genes.

**S4 Table. Enriched GO terms in the GSE39549 microarray data.** Table with the over repressented GO terms using the relevant features found in the GSE39549 microarray data and DAVID. The table has the GO term, its description, the p-value, adjusted p-value, q-value and its relevant genes. The rows highlighted in yellow are related to immune response.

**S5 Table. Relevant features found in the TGP microarray data.** Table with the gene symbol of the all relevant features found in the TGP data with the FSHMM strategy.

**S6 Table. Enriched GO terms with in TGP microarray data.** Table with the over repressented GO terms using the relevant features found in the TGP CCl_4_ microarray data and DAVID. The table has the GO term, its description, the p-value, adjusted p-value, q-value and its relevant genes.

**S7 Table. Enriched KEGG pathways in the TGP microarray data.** Table with the over repressented KEGG pathways using the relevant features found in the TGP CCl_4_ microarray data and KEGG. The table has the KEGG pathways, its description, the p-value, adjusted p-value, q-value and its relevant genes.

## Acknowledgments

Fellowship support to RACO by Consejo Nacional de Ciencia y Tecnologia CONACYT, research supported by Tecnológico de Monterrey, Campus Ciudad de México and Instituto Nacional de Medicina Genómica.

## References

1. Jensen R, Shen Q. Computational Intelligence and Feature Selection: Rough and Fuzzy Approaches. IEEE Press. 2007; 1st Edition.

2. Saunders C, Grobelnik M, Gunn S, Shawe-Taylor J. Subspace, Latent Structure and Feature Selection. Springer. 2006; 1st Edition.

3. Guyon I, Elisseeff A. An introduction to variable and feature selection. JMLR. 2003 Mar;3:1157–1182.

4. Liu H, Motoda H. Computational Methods of Feature Selection. CRC Press.; 2007.

5. Chin AJ, Mirzal A, Haron H, et al. Supervised, Unsupervised and Semi-supervised Feature Selection: A Review on Gene Selection. IEEE Transactions on Computational Biology and Bioinformatics. 2015;13(5):971–989.

6. Ma S, Huang J. Penalized feature selection and classification in bioinformatics. Briefings in Bioinformatics. 2008 June;9(5):392–403.

7. Saeys, Y, Inza I, Larranãga P. A review of feature selection techniques in bioinformatics. Bioinformatics. 2007 Oct;23(19):2507–2517.

8. Adams S, Beling P. A survey of feature selection methods for Gaussian mixture models and hidden Markov models. Springer Netherlands.2017:1–41.

9. Jafari P, Azuaje F. An assessment of recently published gene expression data analyses: reporting experimental design and statistical factors. BMC Med. Inform. Decis. Mak. 2006 June;6(27):1–27.

10. Efron B, Tibshirani R, Storey JD, Tusher V, Empirical Bayes analysis of a microarray experiment.. J. Am. Stat. Assoc. 2001 Dec;96(456):1151–1160.

11. Adams S, Beling P, Cogill R. Feature Selection for hidden Markov models and hidden Semi-Markov models. IEEE. Translations and content mining. 2016 April;4(1):1642–1657.

12. Zhu H, He Z, Leung H. Simultaneous Feature and model Selection for Continuous hidden Markov models. IEEE SIGNAL PROCESSING LETTERS. 2012 May;19(5):279–282.

13. Law MHC, Figueiredo MAT, Jain AK. Simultaneous feature selection and clustering using mixture models. IEEE Trans. Patt. Anal. Mach. Intell. 2004 July;26(9):1154–1166.

14. Zheng Y, Jeon B, Sun L, Zhang J, Zhang H. Student’s t-hidden Markov model for Unsupervised Learning Using Localized Feature Selection. IEEE Transactions on Circuits and Systems for Video Technology. 2017 July;9(12):1–10.

15. Barrett T, Wilhite SE, Ledoux P, Evangelista C, et al. NCBI GEO: archive for functional genomics data sets. Nucleic Acids Res. 2013 Jan;41(Database issue):gks119.

16. Uehara T, Ono A, Maruyama T, Kato I, Yamada H, Ohno Y, et al. The Japanese toxicogenomics project: application of toxicogenomics. Molecular nutrition & food research. 2010;54(2):218–227.

17. Hernández-de-Diego R, Boix-Chova N, Gómez-Cabrero D, Tegner J, Abugessaisa I,Conesa A STATegra EMS: an Experiment Management System for complex next-generation omics experiments. BMC Systems Biology. 2014 March;88(Suppl 2):S9.

18. Kwon EY, Shin SK, Cho YY, Jung UJ, Kim E, Park T, YoonPark JH, Yun JN, McGregor RA Time-course microarrays reveal early activation of the immune transcriptome and adipokine dysregulation leads to fibrosis in visceral adipose depots during diet-induced obesity. BMC Genomics. 2012 April;13(1): 450–465.

19. Ferreirós-Vidal I, Carroll T, Taylor B, Terry A, Liang Z, et al. Genome-wide identification of Ikaros targets elucidates its contribution to mouse B-cell lineage specification and pre-B-cell differentiation.. Blood. 2013;;121(10):1769–82.

20. Huang DW, Sherman BT, Lempicki RA. Systematic and integrative analysis of large gene lists using DAVID Bioinformatics Resources. Nature Protoc. 2009;4(1):44–57.

21. Recknagel RO, Glende EA, Dolak JA, Waller RL. Mechanism of Carbon-tetrachloride Toxicity. Pharmacology & Therapeutics. 1989;43 (43): 139–154.

22. Seifert WF, Bosma A, Brouwer A, et al. Vitamin A deficiency potentiates carbon tetrachloride-induced liver fibrosis in rats. Hepatology. 1994;19 (1): 193–201

23. Lee HS, Jung KH, Hong SW, Park IS, Lee C, et al. Morin Protects Acute Liver Damage by Carbon Tetrachloride (CCl_4_) in Rat. Arch Pharm Res. 2008;31(9):1160–1165.

24. Kanehisa M, Furumichi M, Tanabe M, Sato Y, Morishima K. KEGG: new perspectives on genomes, pathways, diseases and drugs. Nucleic Acids Res. 2017;45(D1):D353–D36.

25. Rabiner LR. A Tutorial on hidden Markov models and Selected Applications in Speech Recognition. Proceedings of the IEEE. 1989;77(2): 257–286.

26. Beal M, Ghahramani Z, Rasmussen C. The infinite hidden Markov model. NIPS’01 Proceedings of the 14th International Conference on Neural Information Processing Systems: Natural and Synthetic. Dec 2001;1:577–584.

27. Ibe O. Markov Processes for Stochastic modeling. Oxford.; 2009.

28. Bilmes J. A Gentle Tutorial of the EM Algorithm and its Application to Parameter Estimation for Gaussian Mixture and hidden Markov models International Computer Science Institute.; 1998

29. Dubitzky W, Granzow M, Berrar D. Fundamentals of data mining in genomics and proteomics. Springer Science & Business Media.; 2007.

